# phyloregion: R package for biogeographic regionalization and spatial conservation

**DOI:** 10.1101/2020.02.12.945691

**Authors:** Barnabas H. Daru, Piyal Karunarathne, Klaus Schliep

## Abstract

1. Biogeographical regionalization is the classification of regions in terms of their biotas and is key to understanding biodiversity patterns across the world. Previously, it was only possible to perform analysis of biogeographic regionalization on small datasets, often using tools that are difficult to replicate.
2. Here, we present `phyloregion`, a package for the analysis of biogeographic regionalization and spatial conservation in the R computing environment, tailored for mega phylogenies and macroecological datasets of ever-increasing size and complexity.
3. Compared to available packages, `phyloregion` is three to four orders of magnitude faster and memory efficient for cluster analysis, determining optimal number of clusters, evolutionary distinctiveness of regions, as well as analysis of more standard conservation measures of phylogenetic diversity, phylogenetic endemism, and evolutionary distinctiveness and global endangerment.
4. A case study of zoogeographic regionalization for 9574 species of squamate reptiles (amphisbaenians, lizards, and snakes) across the globe, reveals their evolutionary affinities, using visualization tools that allow rapid identification of patterns and underlying processes with user-friendly colours–for example– indicating the levels of differentiation of the taxa in different regions.
5. Ultimately, `phyloregion` would facilitate rapid biogeographic analyses that accommodates the ongoing mass-production of species occurrence records and phylogenetic datasets at any scale and for any taxonomic group into completely reproducible R workflows.

## 1.0 Introduction

In biogeography, there is growing interest in the analysis of datasets of ever-increasing size and complexity to explain biodiversity patterns and underlying processes. A common approach is biogeographical regionalization, the grouping of organisms based on shared features and how they respond to past or current physical and biological determinants (Kreft & Jetz, 2010; Morrone, 2018). The units of biogeographical regionalization i.e., “phyloregions” or “bioregions”, are key to our understanding of the ecological and historical drivers affecting species distribution in macroecological or large-scale conservation studies (Kreft & Jetz, 2010; Ladle & Whittaker, 2011; Moreno Saiz et al., 2013; Oikonomou et al., 2014; Ficetola et al., 2017; Morrone, 2018). When paired with phylogenetic information, biogeographical regionalization allows geographic regions that do not share any species in common to be quantified (Graham and Fine, 2008), and can identify patterns overlooked by species-level analyses (Holt et al. 2013; Daru et al. 2016). However, compared to the mass-production of species distribution and phylogenetic datasets, statistical and computational approaches necessary to analyze such data, and approaches that can incorporate efficient storage and manipulation of such data, are lacking.

A few open-source tools have recently become available and can provide infrastructural support for analysis of biogeographical regionalization. The *ape* package (Paradis and Schliep 2018) contains a comprehensive collection of tools for analyses of phylogenetics and evolution and is useful for reading, writing, and manipulating phylogenetic trees, among many other functions. The *betapart* package (Baselga & Orme 2012) performs computations of total dissimilarity in species composition along with their respective turnover and nestedness components. *picante* focused on analysis of phylogenetic community structure and trait evolution (Kembel et al. 2010). The use of network methods to detect bioregions (Carstensen and Olesen 2009, Thébault 2013, Vilhena and Antonelli 2015), while not yet implemented in the R computing environment, provides an alternative clustering method based on bipartite networks, and performs well at identifying interzones between regions (Bloomfield et al. 2018). However, there is no consensus on which method is the most appropriate for biogeographical regionalization and spatial conservation at large scales (Dapporto et al., 2015; Bloomfield et al. 2018; Morrone, 2018). The most effective approach to biogeographical regionalization might therefore depend on the system under study and the research questions.

Here, we present `phyloregion` R package that permits the integration of phylogenetic relationships and species distributions for identifying biogeographical regions of different lineages to elucidate the spatial and temporal evolution of biota in a region. Specifically, `phyloregion` provides functions for clustering substantially large-scale species assemblages, determining optimal number of clusters, quantifying evolutionary distinctiveness of phyloregions, and visualizing various facets of alpha and beta (differences in species composition between local communities) diversity. We illustrate the utility of the proposed package with a simulated dataset and an empirical dataset on the flora of southern Africa that includes species distributions and their phylogenetic relationships. Moreover, we also present a case study for zoogeographic regionalization with the most comprehensive dataset on the phylogenetic relationships and geographic distributions for 9574 species of squamate reptiles (amphisbaenians, lizards, and snakes) across the globe, to demonstrate its potential for analysis at any scale and for any taxonomic group. Visualization tools allow rapid identification of phyloregions with colours in multidimensional scaling space indicating levels of differentiation of the taxa in different phyloregions.

## 2.0 Overview and general workflow of phyloregion

The `phyloregion` package interacts with several other R packages including *Matrix* (Bates and Maechler 2019), *ape* (Paradis & Schliep 2018), *betapart* (Baselga & Orme 2012), *raster* (Hijmans 2019), and *sp* (Bivand et al. 2013). We provide a workflow of the `phyloregion` package for biogeographical assessment of any selected taxa and region (**Fig. 1**). The workflow demonstrates steps from preparation of different types of data to visualizing the results of biogeographical regionalization, together with tips on selecting the optimal method for achieving the best output, depending on the types of data used and research questions. The development version of `phyloregion` is hosted on github at https://github.com/darunabas/phyloregion. To install `phyloregion`, in R, type:

**Fig. 1.**
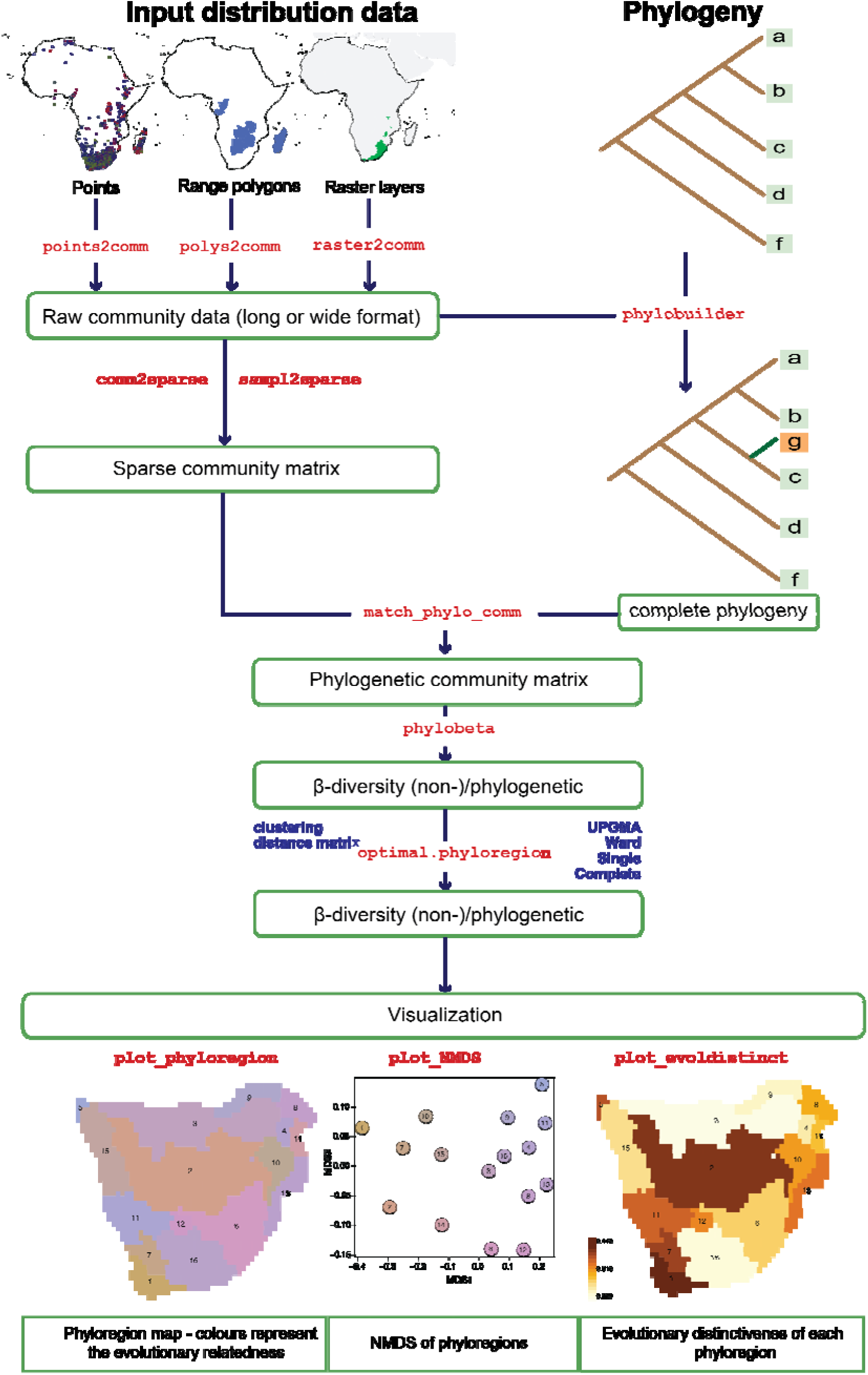
Typical workflow for analysis of biogeographical regionalization using `phyloregion`. a) Distribution data (point records, polygons, and raster layers) is converted to a long community data frame format before b) conversion to a sparse community matrix. When paired with phylogenetic data, `phylobuilder` creates a subtree with largest overlap from a species list, thereby ensuring complete representation of missing data. c) phylocommunity matrix to visualization of results.

~~~
if (!requireNamespace(“devtools”, quietly = TRUE))
   install.packages(“devtools”)
devtools::install_github(“darunabas/phyloregion”)
library(phyloregion)
~~~

### 2.1. Raw Data

#### 2.1.1 Distribution data input

The `phyloregion` package ships with functions for manipulating at least three categories of distribution data at varying spatial grains and extents: point records, extent-of-occurrence polygons and raster layers. Extent-of-occurrence range maps can be derived from the IUCN Redlist spatial database (https://www.iucnredlist.org/resources/spatial-data-download), published monographs or field guides validated by taxonomic experts. Point records are commonly derived from GBIF, iDigBio, or CIESIN and typically have columns of geographic coordinates for each observation. Raster layers are typically derived from analysis of species distribution modeling, such as *aquamaps* (Kaschner et al. 2016). An overview can be easily obtained with the functions `points2comm, polys2comm` and `raster2comm` for point records, polygons, or raster layers, respectively. Depending on the data source, all three functions ultimately provide convenient interfaces to convert the distribution data to a community data frame at varying spatial grains and extents for downstream analyses.

#### 2.1.2 Phylogenetic data

Phylogenies are often derived from DNA sequences or supertree approaches, however, they tend to be prevalent with missing taxa for most non-charismatic groups e.g. plants or insects. When paired with distribution data, phylogenies can aid the discovery of common patterns and processes that underlie the formation of biogeographic regions (Wiley 1988, Daru et al. 2017). The function `phylobuilder` appends missing taxa to a supertree. Unlike other tree-building algorithms that manually graft missing taxa into a working supertree, `phylobuilder` creates a subtree with the largest overlap from a species list at a fast speed. If species in the taxon list are not in the tree (tip label), species will be added at the most recent common ancestor at the genus or family level when possible.

## 3.0 Data preparation and analyses

### 3.1. Sparse community matrix

A community composition dataset is commonly represented as a matrix of 1s and 0s with species as columns and rows as spatial cells or communities. In practice, such a matrix can contain many zero values because species are known to generally have unimodal distributions along environmental gradients (ter Braak & Prentice, 1988), and storing and analyzing every single element of that matrix can be computationally challenging and expensive. Indeed, for large matrices, most base R functions cannot make a table with > 2^31 elements. One approach to overcome this limitation is to utilize sparse matrix, a matrix with a high proportion of zero entries (Duff 1977). Because a sparse matrix is comprised mostly of 0s, it only stores the non-zero entries, from which several measures of biodiversity including biogeographical regionalization can be calculated. Our `sampl2sparse` function allows conversion of community data from either long or wide formats to a condensed sparse matrix (**Fig. 2**) to ease downstream analyses such as compositional dissimilarity and avoid the exhaustion of computer memory capacities.

**Fig. 2.**
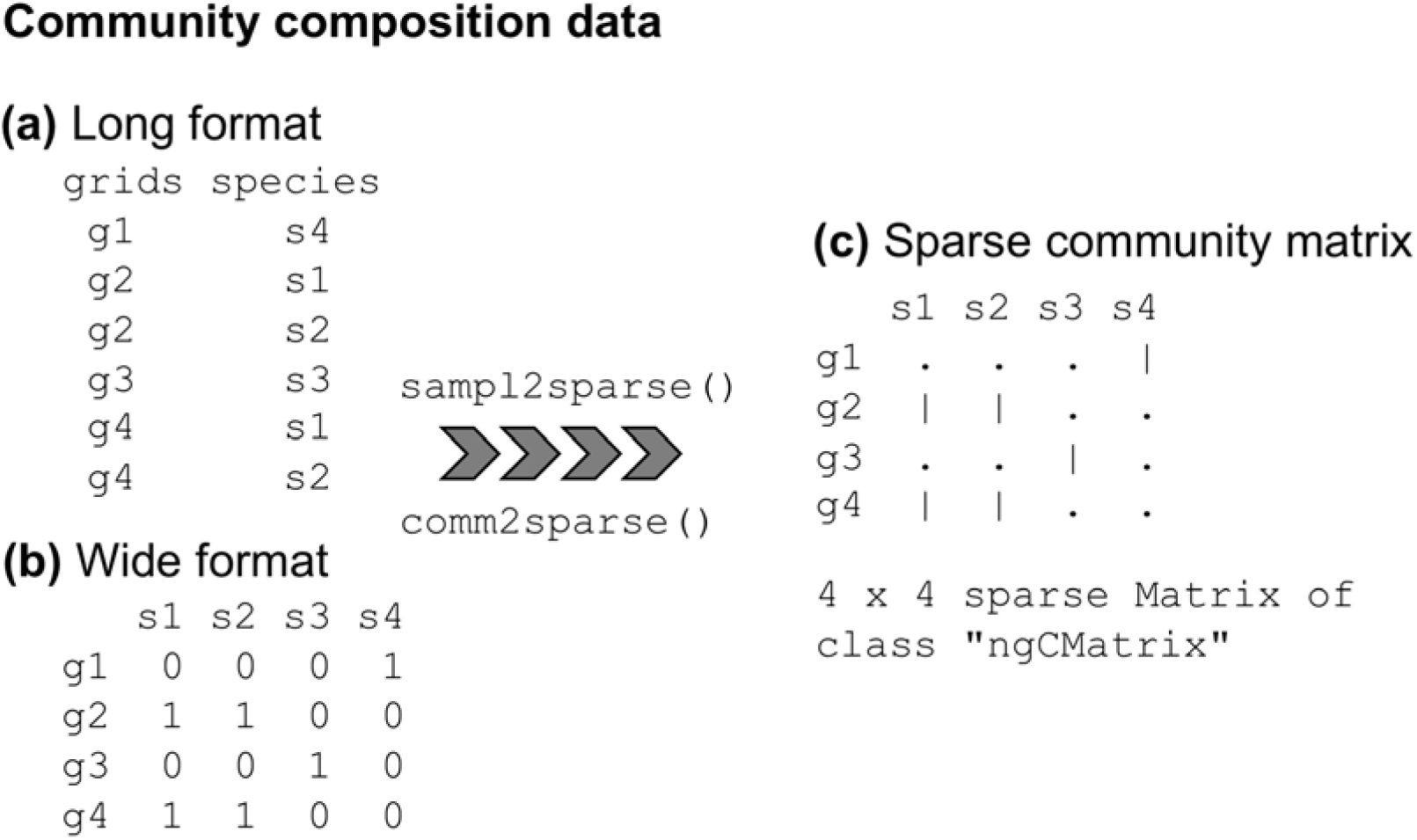
Illustration showing community data conversion to sparse community matrix by (a) `sampl2sparse` function when the raw data is in long community data format, or (b) `com2sparse` for wide community data format. The result is (c) a sparse community matrix for downstream analysis.

### 3.2. Matching phylogeny and community composition data

In community ecology and biogeographic analyses, it is sometimes desirable to make sure that different datasets (e.g. community, phylogeny and trait) match one another (Kembel et al. 2010). However, existing tools are not tailored for comparing taxa in mega phylogenies spanning thousands of taxa with community composition datasets at large scales. We present `match_phylo_comm` that compares a sparse community matrix against a phylogenetic tree and adds missing species to the tree at the genus or higher taxonomic levels.

### 3.3. Generating beta diversity (phylogenetic and non-phylogenetic)

The three commonly used methods for quantifying β-diversity, the variation in species composition among sites, – Simpson, Sorenson and Jaccard (Laffan et al. 2016) – are included in the package as a comparative and optimal selection tool. The `phyloregion`’s functions `beta_diss` and `phylobeta` compute efficiently pairwise dissimilarities matrices for large sparse community matrices and phylogenetic trees for taxonomic and phylogenetic turnover, respectively. The results are stored as distance objects for later use.

### 3.4. Cluster algorithm selection and validation

To overcome the lack of *a priori* justification for using a particular method for identifying phyloregions, the function `select_linkage` can contrast nine widely used hierarchical clustering algorithms (including UPGMA, and single linkage) on the (phylogenetic) beta diversity matrix for degree of data distortion using Sokal & Rohlf’s (1962) cophenetic correlation coefficient. The cophenetic correlation coefficient measures how faithfully the original pairwise distance matrix is represented by the dendrogram (Sokal & Rohlf, 1962). Thus, the best method is indicated by higher correlation values, resulting in regions with a maximum internal similarity but with maximum differences from other regions.

### 3.5. Determining the optimal number of clusters

The function `optimal.phyloregion` utilizes the efficiency of the so-called “elbow” (also “knee”) method corresponding to the point of maximum curvature (Salvador & Chan, 2004), to determine the optimal number of clusters that best describes the observed (phylogenetic) beta diversity matrix. Depending on the research question, the scale of the cutting depth or clustering algorithm `method` can be varied systematically. The output is used to visualize relationships among phyloregions using hierarchical dendrograms of dissimilarity and NMDS ordination, and are assessed for spatial coherence by mapping and/or quantifying their evolutionary distinctiveness.

### 3.6. Evolutionary distinctiveness of phyloregions

The function `ed_phyloregion` estimates evolutionary distinctiveness of each phyloregion by computing the mean value of (phylogenetic) beta diversity between a focal phyloregion and all other phyloregions in the study area. It takes a distance matrix and returns a “phyloregion” object containing a phyloregion × phyloregion distance object. Areas of high evolutionary distinctiveness can provide new insights in the mechanisms that are responsible for generating ecological diversity such as speciation, niche conservatism, extinction and dispersal (Holt et al. 2013; Daru et al. 2017).

## 4.0. Visual representation and assessment of biogeographic regions

The `phyloregion` package also provides a number of functions that aid elaborate visualization and assessment of biogeographic regions.

- `plot_phyloregion` can display clusters of cells (i.e. ‘phyloregions’ or ‘bioregions’) in multidimensional scaling colour space matching the colour vision of the human visual system (Kruskal 1964). The colours indicate the levels of differentiation of clades in different phyloregions. Phyloregions with similar colours have similar clades and those with different colours differ in the clades they enclose (**Fig. 1**).
- `plot_evoldistinct` quantifies evolutionary distinctiveness of phyloregions in geographic space as the mean of pairwise beta diversity values between each phyloregion and all other phyloregions and displays them in HCL colour space (default is “`YlOrBr`” palette; **Fig. 1**). Darker regions indicate regions of higher evolutionary distinctiveness.
- `plot_swatch` maps discretized values of a quantity based on continuous numerical variables of their cells or sites for visualization as heatmap in sequential colour palettes.

## 5.0 Case study of biogeographical regionalization of squamate reptiles

We validated the application of the `phyloregion` package on the geographic distributions and phylogenetic data for all 9574 species of squamate reptiles across the globe (data: Tonini et al. 2016). Despite the fact that reptiles were part of the dataset used in Wallace’s original zoogeographic regionalization along with birds, mammals and insects (Wallace 1876), they have been largely neglected in modern regionalization schemes (Kreft & Jetz 2010; Holt et al., 2013; Meiri & Chapple 2016). Nevertheless, squamate reptiles are one of the most diverse and widely distributed animal groups in the world (Böhm et al. 2013). Most notably, due to the high extinction rates they are facing, the distribution data, phylogeny, and evolutionary relatedness of squamates have recently been well documented (Tonini et al. 2016 and references therein). These make squamate reptiles an ideal system to test the robustness and implementation of `phyloregion` for biogeographic regionalization and spatial conservation at large scales.

We used updated extent-of-occurrence polygons representing the maximum geographical extent of each squamate reptile species (Roll et al. 2017). We ran the `polys2comm, sampl2sparse,` and `match_phylo_comm` wrapper functions to generate the community data at a resolution of 1°× 1°. Note that this resolution can be adjusted by varying the `res` argument in the function `fishnet(mask, res = 0.5)`. We accounted for phylogenetic uncertainty in our analyses by drawing 100 trees at random from a posterior distribution of fully resolved trees (Tonini et al. 2016) to generate phylogenetic dissimilarity matrices (with Simpson’s pairwise phylogenetic dissimilarities as default), and took the mean across grid cells using `mean_matrix`. Note that other dissimilarity indices such as “Jaccard” and “Sorensen” can be used as desired (Laffan et al. 2016), depending on the data used and research questions; review function `phylobeta`.

Using the ‘elbow method’ (function `optimal.phyloregion`), we identified 18 optimal phyloregions (i.e. maximum explained variance of 0.72 for clustering achieved at k = 18) of squamate reptiles (**Fig. 3**). UPGMA was identified as the best clustering algorithm (cophenetic correlation coefficient = 0.8; selected using function `select_linkage`).

**Fig. 3.**
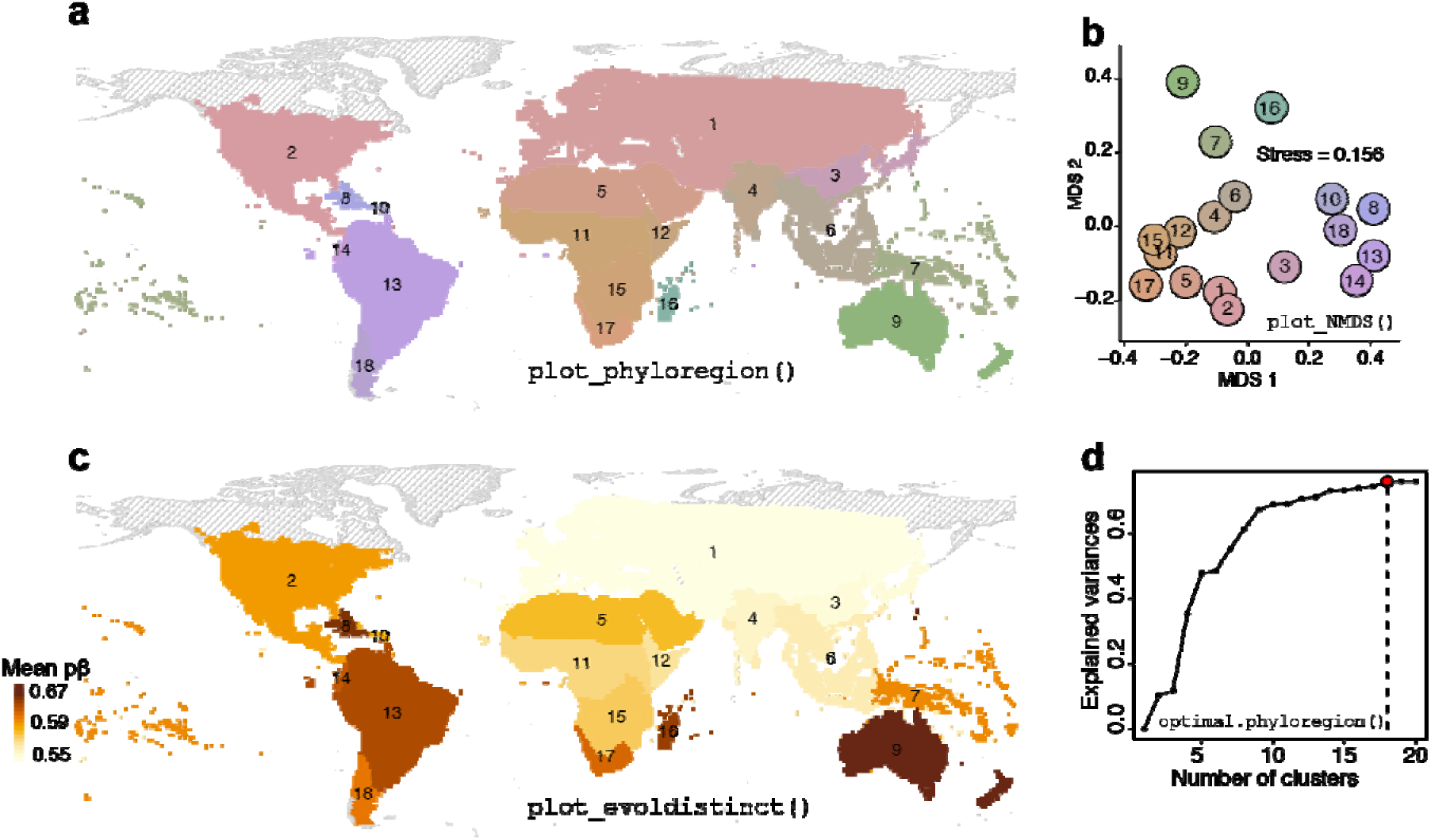
A global phylogenetic regionalization of 9574 species of squamate reptiles reveals their evolutionary affinities. **a**, Map of phyloregions shows evolutionary affinities among disjunct assemblages (function `plot_phyloregion`). **b**, The ordination of phyloregions in NMDS space shows a tropical-temperate divide (function `plot_NMDS`). **c**, evolutionary distinctiveness is high in the tropics than temperate bioregions (function `plot_evoldistinct`). **d**, the optimal number of phyloregions (function `optimal.phyloregion`). Colours differentiating between phyloregions in the map (**a**) and NMDS plot (**b**) are identical.

The resulting phyloregions for squamate reptiles show substantial congruence to Holt et al.’s (2013) updates of Wallace’s original zoogeographic regions including Oceanian, Australian, Madagascan, Palearctic and Nearctic (**Fig. 3a**). However, we also identified some discrepancies. For example, the Afrotropical realm (*sensu* Holt et al. 2013) was divided into four phyloregions in our study corresponding to West and Central Africa (11), Horn of Africa (12), Zambezian (15), and South African (17). We also identified a new phyloregion overlapping Chile-Patagonian in temperate South America. This discrepancy might be due to the focal group being reptiles whereas Holt et al. present results for birds, mammals and amphibians; or differences in spatial grain size (1°× 1° in our study vs 2°× 2° in Holt et al. (2013)). Phylogenetic beta diversity and environmental correlates are systematically grain (spatial resolution) dependent (e.g. Keil et al. 2012).

Notably, geographically proximal phyloregions tend to have low levels of faunal similarity (**Fig. 3b**), suggesting spatial patterns of species diversity can have different phylogenetic structures (Hawkins et al. 2012). Mean phylogenetic turnover of squamate reptiles between a phyloregion and all other phyloregions (function `ed_phyloregion`) indicates a latitudinal gradient in evolutionary distinctiveness, with higher evolutionary distinctiveness in the tropics than in temperate phyloregions (**Fig. 3c**), a similar observation to Tonini et al. (2016). Notably, the Australian phyloregion has the highest mean phylogenetic turnover (mean phylogenetic turnover between Australian and all other phyloregions = 0.67; **Fig. 3c**), reflecting strong niche conservatism or limited dispersal of lineages in this phyloregion.

The use of phylogenetic information and species distributions allows a deeper understanding of the mechanisms determining current patterns of biodiversity. Our evolutionary distinctiveness analysis in the recognized phyloregions brings a new component of evolutionary importance of each region to the biogeographic regionalization as well as for conservation prioritization. Most of the phyloregions found here spanned multiple ecoregions and biogeographic realms, suggesting that conservation planning should be adjusted to cover these larger phyloregions.

## 6.0. phyloregion as tool for spatial conservation

We demonstrate the utility of `phyloregion` in mapping standard conservation metrics of species richness, weighted endemism (`weighted.endemism`) and threat (`mapTraits`) as well as fast computations of phylodiversity measures such as phylogenetic diversity (`PD`), phylogenetic endemism (`PE`), and evolutionary distinctiveness and global endangerment (`EDGE`). The major advantage of these functions compared to available tools e.g. biodiverse (Laffan et al. 2010), is the ability to utilize sparse matrix that speeds up the analyses without exhausting computer memories, making it ideal for handling any data from small local scales to large regional and global scales.

## 6.0. Benchmarking phyloregion

We compared the execution time of `phyloregion`’s functions with available packages using exactly the same datasets (R code for benchmarking `phyloregion` with available packages is provided as Data S1). Regardless of the size of the distribution data and phylogenetic tree, `phyloregion` is 3 or 4 orders of magnitude faster and memory efficient (**Fig. 4**).

**Fig. 4.**
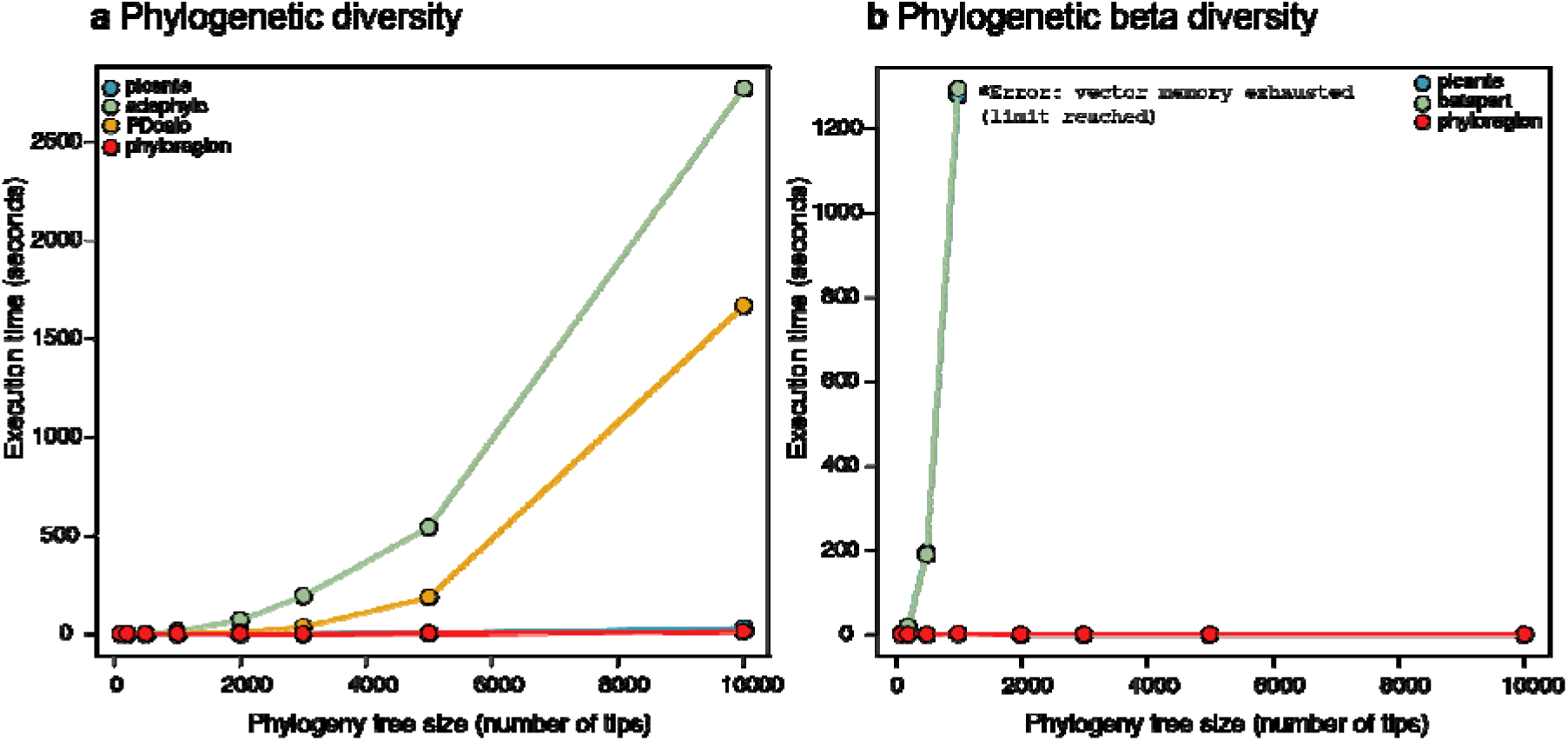
Benchmarking `phyloregion` with available packages in analysis of **a**, phylogenetic diversity, and **b**, phylogenetic beta diversity.

## 7.0. Concluding Remarks

Despite the few other tools that have provided support for biogeographic regionalization and spatial conservation e.g. *ape* (Paradis & Schliep 2018), betapart (Baselga & Orme 2012), or *vegan* (Oksanen et al. 2019) among many others, `phyloregion` adds the following novelties compared to available packages: 1) ability to utilize sparse matrix and large-scale phylogenies for analysis of biogeographical regionalization and spatial conservation, allowing normal operations of a typical matrix in base R to be done on the sparse matrix, 2) novel functions for speedy raw data conversion to sparse community matrix as well as a user-friendly analysis of biogeographical regionalization into completely reproducible R workflows, 3) although the functionality of the package has been developed with biogeographical regionalization in mind, it can accommodate analysis of spatial conservation at large scales such as mapping various biodiversity metrics for conservation ranging from mapping biodiversity hotspots of species richness, endemism, or threat. Other implementations of `phyloregion` include the addition of phylogenetic information and sparse community matrix to map evolutionary diversity including phylogenetic diversity, phylogenetic endemism, and evolutionary distinctiveness and global endangerment.

Overall, no hard rule exists on how to perform analysis of biogeographic regionalization or spatial conservation - the choice of approach will ultimately depend on the goal of the study, questions, hypotheses or the taxonomic group. The goal of `phyloregion` is to facilitate analysis of biogeographic regionalization and spatial conservation at any scale and for any taxonomic group, tailored to accommodate the ongoing mass-production of species occurrence data and phylogenetic datasets.

## Acknowledgements

B.H.D. is supported by Texas A&M University at Corpus Christi.

## Authors’ contributions

B.H.D. conceived the project. B.H.D. and K.S. developed the method. B.H.D., K.S., and P.K. tested the method. B.H.D. analyzed the data and led the writing with help from P.K. All co-authors assisted with edits and approve publication.

## Data accessibility

The `phyloregion` R package and documentation are hosted at https://github.com/darunabas/phyloregion. All data and scripts necessary to repeat the analyses for the squamate reptiles described here have been made available through the Dryad Digital Data Repository https://doi.org/10.5061/dryad.tdz08kpw6 (Daru et al. 2019).

